# DNA polymerase beta expression in head & neck cancer modulates the poly(ADP-ribose)-mediated replication checkpoint

**DOI:** 10.1101/2025.01.15.633225

**Authors:** Md Maruf Khan, Wynand P. Roos, Anusha Angajala, Denise Y. Gibbs, Jeffrey C. Liu, Camille Ragin, Robert W. Sobol

## Abstract

Head and Neck Squamous Cell Carcinoma (HNSCC) imposes a significant health burden, necessitating innovative therapeutic strategies to enhance treatment efficacy. Current treatments such as surgery, radiation, and chemotherapy have limited effectiveness and yield severe side effects, emphasizing the need for targeted therapies. We have focused on DNA polymerase beta (Polβ) and its roles in replication stress, cellular responses to DNA damaging therapies, and DNA damage response modifiers. Our investigations reveal a regulatory role for base excision repair (BER) proteins, including Polβ, in the cellular response to inhibitors of poly(ADP-ribose) glycohydrolase (PARG), an enzyme involved in poly(ADP-ribose) (PAR) degradation. The inhibition of PARG, in HNSCC cells, elicits replication stress. Further, this activates the PAR-induced S-phase/ATR checkpoint, leading to a block to replication, cell cycle arrest, and the onset of apoptosis. However, Polβ overexpression mitigates this response, reducing replication-stress induced PAR foci formation, suggesting a modulation of replication checkpoint activation. We found that PARG inhibitor treatment is ineffective on HNSCC cells that overexpress Polβ, implying that the PARG inhibitor-induced PAR and apoptotic response is dependent on the level of Polβ. Further, our *in vitro* experiments demonstrate that combining PARG and ATR/CHK1 inhibitors overcomes Polβ-mediated treatment resistance in HNSCC cells, producing enhanced effects as compared to the individual treatment conditions. Our findings suggest a possible treatment paradigm for HNSCC, employing ATR or CHK1 inhibitors in combination with PARG inhibitors. This strategy offers a promising path for more effective HNSCC treatments, potentially overcoming Polβ-related resistance.

## 1. Introduction

Head and Neck Squamous Cell Carcinoma (HNSCC) stands as a formidable health challenge, necessitating the exploration of innovative therapeutic strategies to augment treatment efficacy [1]. Current standard interventions, comprising surgical intervention, radiation therapy, and chemotherapy, are frequently constrained by their limited effectiveness and the induction of substantial side effects. This underscores an imperative for novel therapeutic modalities that selectively target cancerous cells while preserving normal tissue integrity [2]. In HNSCC, it has been suggested that overexpression of DNA polymerase beta (Polβ) may be related to genetic ancestry [3, 4]. In a previous study, altered mRNA expression of Polβ was found to be associated with ancestry-informative markers (AIMs), suggesting elevated levels of Polβ among African American (AfAm) patients compared to their European American (EuAm) counterparts that are further associated with poor survival response following therapy [4].

Our investigative focus was directed towards comprehending the intricate involvement of Polβ in the context of replication stress, response to DNA damaging therapies, and the modulation of DNA damage response modifiers in HNSCC. Given the critical role of Polβ in DNA repair [5, 6], it is suggested that elevated Polβ expression may engender resistance to some therapeutic approaches in this malignancy. Notably, our inquiries unveiled a regulatory role for base excision repair (BER) proteins, particularly Polβ, in the cellular response to inhibitors of poly(ADP-ribose) glycohydrolase (PARG), a pivotal enzyme involved in the degradation of poly(ADP-ribose) (PAR) [7].

Poly(ADP-ribose)ylation (PARylation) catalyzed by poly(ADP-ribose) polymerase 1 (PARP1) and its subsequent reversal by PARG orchestrates an intricate DNA damage response [8]. Maintaining PAR homeostasis is pivotal in regulating these responses by dynamically modulating protein-protein and protein–DNA interactions crucial for genomic stability and cellular viability [9–14]. Inhibition of PAR synthesis via PARP inhibitors (PARPi) disrupts signaling pathways and DNA repair mechanisms, resulting in defects in replication and enhanced cell death depending on the genetic background [15]. Conversely, PARG inhibitors [16] cause delayed repair of both single- and double-strand DNA breaks and a defect in polymerase theta-mediated end-joining [17]. This is accompanied by aberrant cell cycle progression [18] and enhanced cell death that appears to depend on the accumulation of replication gaps [19] and the level of NAD^+^ [18]. PARG, encoded by a single gene with alternative splicing generating five isoforms, functions as an endo- and exo-glycohydrolase, swiftly degrading PAR synthesized by PARP1 to coordinate DNA repair processes [13]. Through hydrolysis of α(1′′-2′) O-glycosidic linkages within PAR chains, PARG liberates ADP-ribose and oligo-(ADP-ribose) chains, which may serve as signaling molecules in response to genotoxic stress [20].

The inhibition of PARG, as elucidated in our previous investigations, elicits a robust intra-S-phase response, activating the ATR and CHK1 signaling pathways [18, 21–23]. Our *in vitro* studies herein delineate a nuanced relationship, where response to the combination of an ATR inhibitor and a PARG inhibitor is intricately linked to Polβ expression in HNSCC cells. The enhance cell-killing effects observed in this combinatorial approach surpass the efficacy exhibited by the individual inhibitors [24]. In parallel, the efficacy of the combination of CHK1 and PARG inhibition is equally effective in HNSCC cells, as also demonstrated in ovarian cancer cells [21], suggesting a potent dual therapeutic strategy. These promising combinations hold considerable potential for the treatment of HNSCC, particularly in instances characterized by heightened Polβ expression [4].

## 2. Materials and methods

### 2.1. Chemicals and reagents

DMEM and RPMI1640 were from Corning (Manassas, VA). MEM, L-glutamine, hygromycin, and penicillin/streptomycin were from Gibco (NY, USA). PDD00017273 (PARG inhibitor; Sigma-Aldrich), AZD6738 (ATR inhibitor; Selleckchem), and MK8776 (CHK1 inhibitor; Selleckchem) were dissolved in DMSO to prepare a stock solution at a concentration of 100mM and stored at −30°C. All plasmids, chemicals, and other reagents used in this study are listed in **Table S1**.

### 2.2. Cell lines and cell culture conditions

293-FT cells were from Thermo Fisher Scientific, FaDu cells were from ATCC, and JHU029 cells were a generous gift from David Sidransky and Jeffrey N. Myers [25]. 293-FT cells were cultured in DMEM (Gibco, Cat#10-013-CV) supplemented with 10% Heat-inactivated (HI) FBS (Bio-Techne, Cat#S11150H) and 1% Penicillin-Streptomycin (Gibco, Cat# 15140122). FaDu cells were grown in MEM (Gibco, Cat# 11095-080) supplemented with 10% HI FBS (Atlanta Biologics, Cat# S11150), and 1% Penicillin-Streptomycin (Gibco, Cat# 15140122). JHU029 cells were grown in RPMI (Corning, Cat# 10-040-CV) supplemented with 10% HI FBS (Atlanta Biologics, Cat# S11150), L-glutamine (Gibco, Cat# 25030-08), sodium pyruvate (Invitrogen, Cat# 11360-070), non-essential amino acids (Invitrogen, Cat# 11140-050) and 1% Penicillin-Streptomycin (Gibco, Cat# 15140122). All cells were grown in a humidified incubator at 37°C with 5% CO_2_. We routinely validate cell lines by Labcorp cell line authentication (recently tested January 2025).

### 2.3. Antibodies

The rabbit anti-XRCC1 antibody was from Bethyl (Cat# A300-065A, 1:2000). The anti-phospho-H2AX (Ser139) antibody was from Cell Signaling (#9718S, 1:1000), the rabbit anti-Histone H3 antibody was from Active Motif (Cat# 39451, 1:4000), and the rabbit anti-DNA Polβ antibody was from Abcam (Cat# ab175197, 1:2000). The goat anti-rabbit IgG-HRP antibody was from Bio-Rad (Cat# 1706515, 1:2000), and the Alexa Fluor 568 goat anti-rabbit antibody was from Invitrogen (Cat# A11011, 1:500).

### 2.4. Plasmid and vector development

Lentiviral vectors were designed in-house and synthesized by Vectorbuilder Inc: pLV-Hygro-EF1a-myc-Polβ(PAMmut) (also available at Addgene, Cat#176150) encodes both a Myc tag fused to the N-terminus of Polβ and a hygromycin resistance cassette [10]. The LivePAR vector, pLV-EF1A-LivePAR-Hygro (also available at Addgene, Cat#176063), expresses a PAR binding domain (aa 100-182 from RNF146) fused to EGFP and also encodes a hygromycin resistance cassette [10, 26, 27].

### 2.5. Lentivirus production and cell transduction

Lentiviral particles were generated by co-transfection of lentiviral packaging and shuttle vectors. A culture of 293-FT cells (1×10^6^/dish) was seeded in a 60mm dish and subjected to an overnight incubation period. The third-generation packaging vectors, namely pMDLg/pRRE (Addgene, Cat# 12251), pRSV-Rev (Addgene, Cat# 12253), and pMD2.G (Addgene, Cat# 12259), alongside the shuttle vector/transfer vector, were co-transfected into 293-FT cells utilizing the TransIT-X2 Dynamic Delivery System (Cat# MIR 6003). Following a 48-hour incubation, the supernatant containing the lentivirus was harvested and subsequently filtered through 0.45 μM filters to eliminate cellular debris and isolate viral particles, following established protocols [10, 18].

For target cell preparation, 2×10^5^ cells per well were seeded into a six-well plate and allowed to grow for 24 hours. Subsequently, the culture medium was replaced with 1 mL of media containing polybrene (2 μg/ml), followed by the gradual addition (dropwise) of the lentiviral particle solution (1 mL). Following an overnight incubation at 32°C, the lentivirus-containing media was replaced with fresh media, and cells were allowed to grow to 70-80% confluency at 37°C. For stable cell lines, cells were cultured for 48 hours at 37°C followed by selection with antibiotics (hygromycin: FaDu, 250 μg/ml; JHU029, 75 μg/ml) for 1–2 weeks [26].

### 2.6. Cell lines with targeted CRISPR/Cas9-mediated knockout of the POLB gene

We developed FaDu and JHU029 cell lines with a stable knockout of POLB using the CRISPR/Cas9 system [10, 18]. FaDu/Polβ-KO and JHU029/Polβ-KO cellular models were established by introducing non-viral ribonucleoprotein complexes (RNPs) obtained from Synthego. These complexes consisted of purified Cas9 protein and a mixture of three single-guide RNAs (gRNA#1 GGCCGCCAUGAGCAAACGGA, gRNA#2 GUGCUAACCUGUGAGCAUGU, gRNA#3 GUCGGUGUGGGAGAGAAGGA), specifically designed to target exon 1 of the POLΒ gene.

FaDu or JHU029 cells were seeded at a density of 2×10^5^ cells/well in a six-well plate and allowed to undergo a 24-hour incubation period. The preparation of the ribonucleoprotein transfection complex involved combining the three single-guide RNAs (sgRNAs), purified Cas9 protein, and the CRISPRMAX-Cas9 transfection reagent (Thermo Fisher Scientific, Cat# CMAX00008) in serum-free OptiMEM (Thermo Fisher Scientific, Cat# 31985070). Following a 30-minute incubation, the transfection complexes were carefully added dropwise to the cells. After 2-3 days, the cells of this knockout (KO) pool were harvested by trypsinization. To isolate single-cell clones, 150 cells from the KO pool were seeded into a 100mm dish, and subsequently, single-cell clones were isolated and expanded. The validation of Polβ knockout (Polβ-KO) was achieved through immunoblot analysis, utilizing a Polβ antibody (Abcam, Cat# ab175197), to confirm the absence of Polβ protein expression. Furthermore, H3 protein expression was evaluated using an H3 antibody (Active Motif, Cat# 39451) to ensure uniform protein loading.

### 2.7. Cell protein extract preparation

Protein extracts, specifically whole-cell lysates (WCL), were generated from cells subjected to various genetic modifications and/or exposed to drugs for varying durations, as delineated in the text. Protein extract of parental and genetically modified human cancer cells was prepared using 2X clear Laemmli buffer (2% SDS, 20% glycerol, 62.5mmol/l Tris-HCl pH 6.8) as we described previously [10, 28]. Cells were initially seeded into a 60 mm cell culture dish and then cultured at 37°C, 5% CO2. Upon reaching 60%–70% confluency, cells underwent two washes with cold phosphate-buffered saline (PBS), after which they were harvested and lysed using an appropriate volume of 2x clear Laemmli buffer. Subsequently, the cell lysates were subjected to a 10-minute boiling step and quantified using the NanoDrop 2000c Spectrophotometer [29].

### 2.8. Immunoblot analysis

Whole-cell lysates (40-50 μg protein) were applied to precast NuPAGE® Novex® 4– 12% Bis-Tris gels and electrophoresed for 1 hour at 100-120V (1.5 hours), as described previously [18]. Following electrophoresis, the separated proteins were transferred to a nitrocellulose membrane using a Turboblotter (Bio-Rad). The membrane was initially blocked with B-TBST (TBS buffer containing 0.05% Tween-20 and supplemented with 5% blotting-grade non-fat dry milk; Bio-Rad) for 1 hour at room temperature. Subsequently, the membrane was incubated overnight at 4°C with primary antibodies in B-TBST. **Table S1** provides details on the primary antibodies and their respective dilutions. After washing, the membranes were treated with secondary antibodies in B-TBST for 1 hour at room temperature. The secondary antibodies used were Bio-Rad goat anti-mouse-HRP conjugate or Bio-Rad anti-rabbit-HRP conjugate. Following additional washing steps, the membrane was exposed to a chemiluminescent substrate, and protein bands were visualized using a Bio-Rad Chemi-Doc MP imaging system.

### 2.9. Detecting poly(ADP-ribose) (PAR) foci upon PARG inhibitor treatment

Poly(ADP-ribose) (PAR) foci analysis was performed using the genetically encoded LivePAR probe [10, 26, 27]. FaDu cells (derived from a EuAm HNSCC patient) and JHU029 cells (derived from a AfAm HNSCC patient) expressing the LivePAR probe were seeded at a density of 2.2 × 10⁵ cells per 60 mm dish containing sterile coverslips. The cells were incubated for 32-36 hours at 37°C in a humidified atmosphere with 5% CO₂ to ensure optimal cell adhesion and growth. Following incubation, cells were treated with media containing 0.1% DMSO (control), or PARG inhibitor (5µM, PDD00017273) at 37°C with 5% CO₂ for 8 hours. The media was then removed, and the cells were washed three times with PBS and fixed with 4% formaldehyde for 15 minutes at room temperature. Permeabilization was achieved by incubating the cells in a 1:1 mixture of methanol and acetone for 10 minutes at −20°C. Coverslips were subsequently washed with PBS and mounted onto glass slides using Vectashield mounting medium. Micrographs were acquired using a Nikon Ti2-E inverted confocal microscope with Ax-R equipped with Ax-R 2k Resonant + Galvo Scan Head. ImageJ software (NIH) was used to quantify the number and intensity of PAR foci across different treatment groups. Data were analyzed to assess the impact of PARG inhibition on PAR foci formation.

### 2.10. Apoptosis analysis

To assess early and late apoptosis, cells were treated with either a DMSO control or PARG inhibitor (5 µM, PDD00017273) for 72 hours. Apoptosis assays were performed using the ViaStain™ No-Wash Annexin V-FITC kit (Nexcelom, CSK-V0007-1) according to the manufacturer’s protocol. The evaluation of apoptotic cells was performed by enumerating cells exhibiting positive signals for annexin-V (detected in the green fluorescence channel; Excitation wavelength: ∼488 nm, Emission wavelength: ∼530 nm) and propidium iodide (detected in the red fluorescence channel; Excitation wavelength: ∼535 nm, Emission wavelength: ∼617 nm) using a Celigo cytometer, employed for the measurement and analysis of fluorescence signals. The quantified signal was normalized against the total cell count, determined by 4’,6-diamidino-2-phenylindole (DAPI) (detected in the blue fluorescence channel; Excitation wavelength: ∼358 nm, Emission wavelength: ∼461 nm).

### 2.11. Induction of phosphorylated-H2AX foci (γH2AX)

FaDu and JHU029 cells were treated with PARGi (5 µM, PDD00017273) and evaluated for the induction of phosphorylated-H2AX (hereafter called γH2AX) foci, essentially as we have described previously [30]. Briefly, coverslips were prepared by immersing them in diethyl ether (FisherSci, Cat# E134–1) for 20 minutes. Following diethyl ether removal, the coverslips were sequentially washed with ethanol (FisherSci, Cat# 2701) at decreasing concentrations (100%, 70%, and ddH₂O). Subsequently, the coverslips were immersed in 1 N HCl (FisherSci, Cat# A144–212) for 20 minutes. Residual HCl was removed by washing the coverslips three times with ddH₂O.

Two coverslips were placed in each 60 mm tissue culture dish, and cells were seeded at a density of 2.0 × 10⁵ cells per dish. After 24 hours, the cells were treated with PARG inhibitor (5 µM, PDD00017273; MilliporeSigma, Cat# SML1781) for an additional 24 hours. Cells were then pre-fixed with 4% formaldehyde in PBS (FisherSci, Cat# J61899.AP) for 15 minutes and subsequently fixed with ice-cold methanol:acetone (7:3) (methanol: FisherSci, Cat# A452SK-4; acetone: FisherSci, Cat# A18–1) at −20°C for 9 minutes. The coverslips were washed in PBS (VWR, Cat# 21–031-CV) and blocked with 5% bovine serum albumin (RPI Research Products International, Cat# A30075–100.0) in 0.25% Triton X-100/PBS (PBST) (Triton X-100: MilliporeSigma, Cat# 93443) for 1 hour at room temperature.

Each coverslip was incubated overnight at 4°C in a humidified chamber with 100 µL of anti-phospho-H2AX (Ser139) antibody (Cell Signaling, Cat# 9718S) diluted 1:1000 in PBST. The coverslips were then washed three times with PBST, followed by incubation with 100 µL of Alexa Fluor 568 goat anti-rabbit antibody (Invitrogen, Cat# A11011) (1:500 in PBST) for 2– 3 hours at room temperature in the dark. After three PBST washes, the coverslips were mounted onto glass slides using Vectashield antifade medium with DAPI (H-1000, FisherSci, Cat# NC1695563) and sealed with clear nail polish (O.P.I. Nature Strong). Micrographs were acquired using a Nikon Ti2-E inverted confocal microscope with Ax-R equipped with Ax-R 2k Resonant + Galvo Scan Head. The number of γH2AX foci per nucleus was determined by using ImageJ software (NIH).

### 2.12. Cell viability assay

Cell viability in response to drug treatments was assessed according to the following protocol [31, 32]. Cells were initially seeded on a 96-well plate at a density of 800 cells per well. After a 24-hour incubation period, cells were exposed to either a single or combined dosage (with multiple dilutions as specified in the figures), without the removal of preconditioned media. Subsequently, after 120 hours (5 days), the total cell population was identified by staining with Hoechst 33342 (2 µM; Thermo Fisher Scientific, Cat# 6224910), and dead cells were identified by staining with propidium iodide (1.5 µM; Sigma, Cat# P4170), followed by a 15-minute incubation at 37°C. Enumeration of both total and dead cells was performed using the Celigo S Image Cytometer (Nexcelom Bioscience, Perkin Elmer). This involved capturing the Hoechst dye signal (excitation/emission wavelength for the blue channel, 377nm/470nm) and the propidium iodide signal (excitation/emission wavelength for the red channel, 531nm/629 nm).

### 2.13. Statistical analysis

Statistical comparisons were performed using Student’s t-test for dose comparisons. Significance was indicated as follows: no stars = not significant, *p < 0.05, **p < 0.01, ***p < 0.001, and ****p < 0.0001. For the LivePAR analysis, statistical significance was determined using a Student’s t-test (****p < 0.0001). For the γ-H2AX analysis, a total of n = 3 independent experiments were performed, with >150 cells analyzed per condition, and statistical significance was assessed using the Mann-Whitney test (two-tailed; ****p < 0.0001). For apoptosis analyses, the Mann-Whitney U test (two-tailed) was used, with n = 3 and >150 samples per condition, and significance was determined at ***p < 0.001, and ****p < 0.0001. All statistical tests were conducted using GraphPad Prism software. Data are presented as mean ± standard deviation (SD).

## 3. Results

### 3.1. Polβ overexpression causes resistance to PARG inhibitors in HNSCC cells

Immunoblot analysis was performed to assess the expression levels of Polβ and XRCC1 in various cell lines, including JHU029, JHU029/Polβ-KO (CRISPR/Cas9 knockout), JHU029/Myc-Polβ (engineered for elevated Polβ expression), FaDu, FaDu/Polβ-KO (CRISPR/Cas9 knockout), and FaDu/Myc-Polβ (engineered for elevated Polβ expression) (**Fig. 1A**). In the JHU029 and FaDu cell lines, the knockout of Polβ (JHU029/Polβ-KO and FaDu/Polβ-KO) resulted in the complete absence of Polβ protein, confirming the successful knockout of the gene. In contrast, overexpression of Polβ (JHU029/Myc-Polβ and FaDu/Myc-Polβ) led to a significant increase in Polβ protein level compared to the parental cell lines, demonstrating effective overexpression. The expression level of XRCC1 remained relatively constant across all cell lines, regardless of Polβ knockout or overexpression status. This indicates that XRCC1 expression is not significantly influenced by the alteration of Polβ expression levels. The consistent expression of Histone H3 (H3) across all samples further supports the validity of the observed results (**Fig. 1A**).

**Fig. 1.**
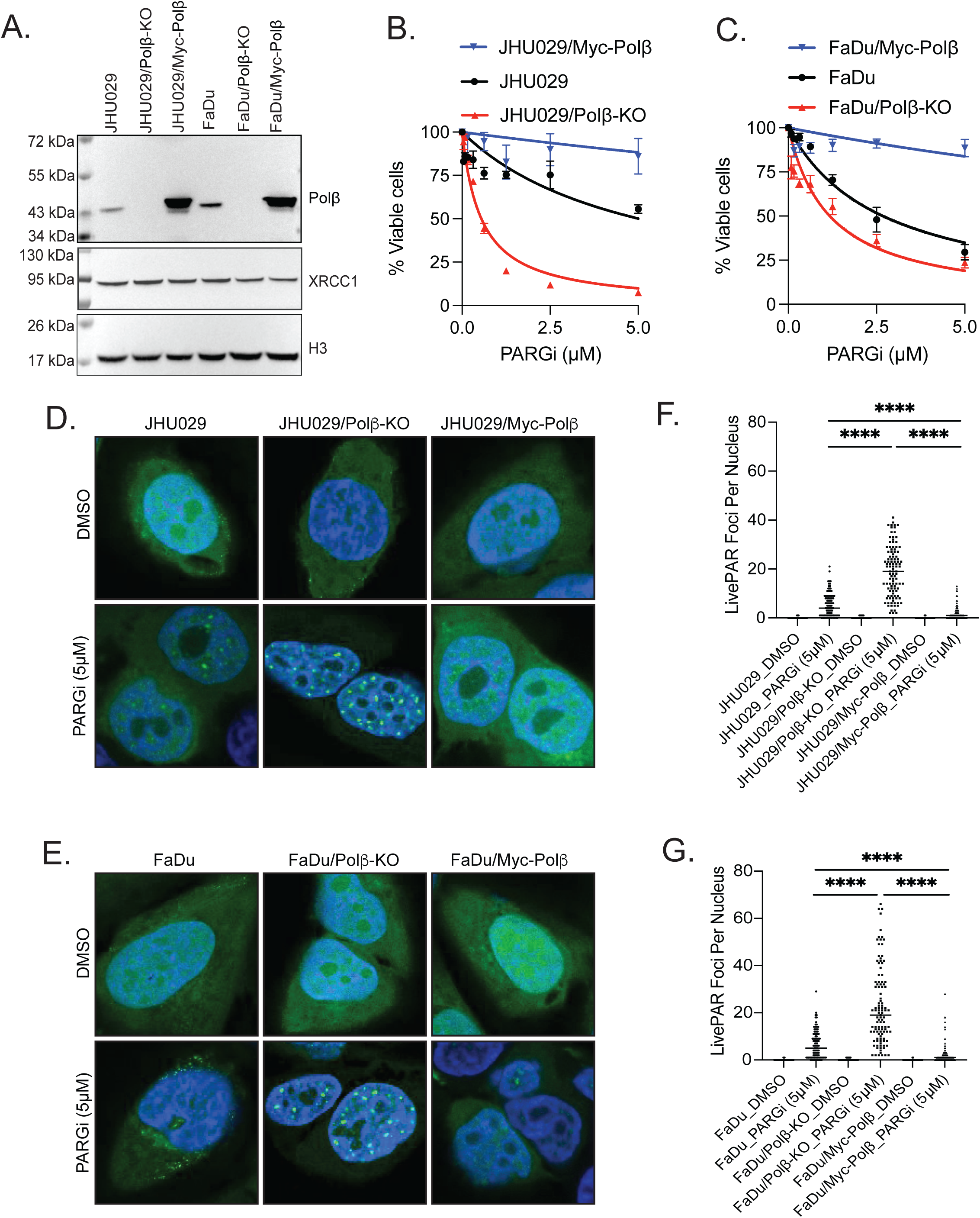
Polβ expression confers resistance to PARG Inhibitors in HNSCC cells. A. Immunoblot analysis of Polβ, XRCC1, and H3 (loading control) in JHU029, JHU029/Polβ-KO, JHU029/Myc-Polβ, FaDu, FaDu/Polβ-KO, and FaDu/Myc-Polβ cell lines. B. Cell viability analysis in JHU029, JHU029/Polβ-KO, and JHU029/Myc-Polβ cells following treatment with the PARG inhibitor, PDD00017273 (0-5 µM) for 120 hours, as indicated, n=3 (comparison of JHU029, JHU029/Polβ-KO or JHU029/Myc-Polβ at each dose: no stars = not significant, p*<0.05, p**<0.01, p***<0.001; Student’s t-test). C. Cell viability analysis in FaDu, FaDu/Polβ-KO, and FaDu/Myc-Polβ cells following treatment with the PARG inhibitor, PDD00017273 (0-5 µM) for 120 hours, as indicated, n=3 (comparison of FaDu, FaDu/Polβ-KO and FaDu/Myc-Polβ at each dose: no stars = not significant, p*<0.05, p**<0.01, p***<0.001; Student’s t-test). D. Fluorescent confocal microscopy images of PAR foci, as detected with the PAR probe (LivePAR), in JHU029, JHU029/Polβ-KO, and JHU029/Myc-Polβ cells treated with 0.1% DMSO (control), or 5 µM PARG inhibitor (PARGi). Distinct PAR foci are visible within the nuclei, indicating increased PAR formation following PARG inhibition for 8 hours. E. Fluorescent confocal microscopy images of PAR foci, as detected with the PAR probe (LivePAR), in FaDu, FaDu/Polβ-KO and FaDu/Myc-Polβ cells treated with 0.1% DMSO (control), or 5 µM PARG inhibitor (PARGi). Distinct PAR foci are visible within the nuclei, indicating increased PAR formation following PARG inhibition for 8 hours. F. Nuclear foci analysis of PAR, as detected with the PAR probe (LivePAR), following exposure of the JHU029, JHU029/Polβ-KO, and JHU029/Myc-Polβ cells to DMSO, or PARGi (5µM) (8 hrs), n=3 (p<****0.001; Student’s t-test). G. Nuclear foci analysis of PAR, as detected with the PAR probe (LivePAR), following exposure of the FaDu, FaDu/Polβ-KO and FaDu/Myc-Polβ cells to DMSO, or PARGi (5µM) (8 hrs), n=3 (p<****0.001; Student’s t-test).

Cytotoxicity analysis demonstrated differential responses to PARG inhibition among the JHU029 cell lines (parental, Polβ-KO, and Polβ overexpression) (**Fig. 1B**). While treatment with the PARG inhibitor (0-5 µM) resulted in a significant decrease in cell viability in the parental and Polβ-KO cells, Polβ overexpression cells exhibited a remarkable resistance to PARG inhibitor-induced cell death. This resistance was evident even at the highest PARG inhibitor concentration, suggesting that elevated levels of Polβ may confer protection against PARG inhibition-mediated cytotoxicity. Similarly, cytotoxicity analysis showed varying responses to PARG inhibition among the FaDu cell lines (parental, Polβ-KO, and Polβ overexpression) (**Fig. 1C**). While PARG inhibitor treatment significantly decreased viability in the parental and Polβ-KO cells, Polβ overexpression cells demonstrated resistance, even at the highest PARGi concentration.

The observed resistance of Polβ overexpression cells to PARG inhibition highlights the complexity of cellular responses to DNA damage and repair mechanisms. It is conceivable that Polβ overexpression may enhance the efficiency of BER [10], or suppress the accumulation of replication-induced single-strand DNA gaps [19], thereby counteracting the cytotoxic effects of PARG inhibition. Alternatively, Polβ overexpression could promote alternative DNA repair pathways or confer survival signals in response to PARG inhibition [33, 34].

### 3.2. Polβ modulates poly(ADP-ribose) (PAR) accumulation and replication checkpoint activation in HNSCC cells following PARG inhibition

Confocal microscopy was utilized to visualize and quantify the formation of PAR foci in the nuclei of JHU029, JHU029/Polβ-KO, and JHU029/Myc-Polβ cells following treatment with PARG inhibitor (**Fig. 1D**). This was also observed when treating FaDu, FaDu/Polβ-KO, and FaDu/Myc-Polβ cells following treatment with PARG inhibitor (**Fig. 1E**). The analysis revealed significant differences in the number of PAR foci among the different cell lines (**Figs. 1F,1G**), dependent on Polβ expression.

FaDu/Polβ-KO cells exhibited a markedly higher number of PAR foci per nucleus as compared to both FaDu and FaDu/Myc-Polβ cells, which displayed relatively low levels of PAR foci. The elevated PAR foci in FaDu/Polβ-KO cells suggest an enhanced activation of the replication checkpoint in the S-phase of the cell cycle, likely due to the accumulation of unrepaired DNA damage or stalled replication forks in the absence of Polβ. Conversely, the reduced PAR foci in FaDu and FaDu/Myc-Polβ cells indicate a more efficient resolution of PARylated intermediates, potentially preventing the activation of the replication checkpoint in response to PARG inhibition.

These findings imply that Polβ plays a crucial role in modulating the cellular response to PARG inhibition, particularly in the context of replication stress, and further underscores the involvement of Polβ in maintaining genome stability during the S-phase of the cell cycle.

### 3.3. γH2AX foci formation in response to PARG inhibition

We then assessed the sensitivity of JHU029, JHU029/Polβ-KO, and JHU029/Polβ overexpression cells to PARG inhibitors by examining the formation of γH2AX foci, a marker of DNA damage and replication stress (**Figs. 2A, S1A**). JHU029 cells exhibited slight resistance to the PARG inhibitor. In contrast, JHU029/Polβ-KO cells showed a marked sensitivity to the PARG inhibitor, with the highest levels of γH2AX foci observed in this cell line. JHU029/Polβ overexpression cells, however, displayed increased resistance to the PARG inhibitor, with the lowest levels of foci formation among the cell lines tested.

**Fig. 2.**
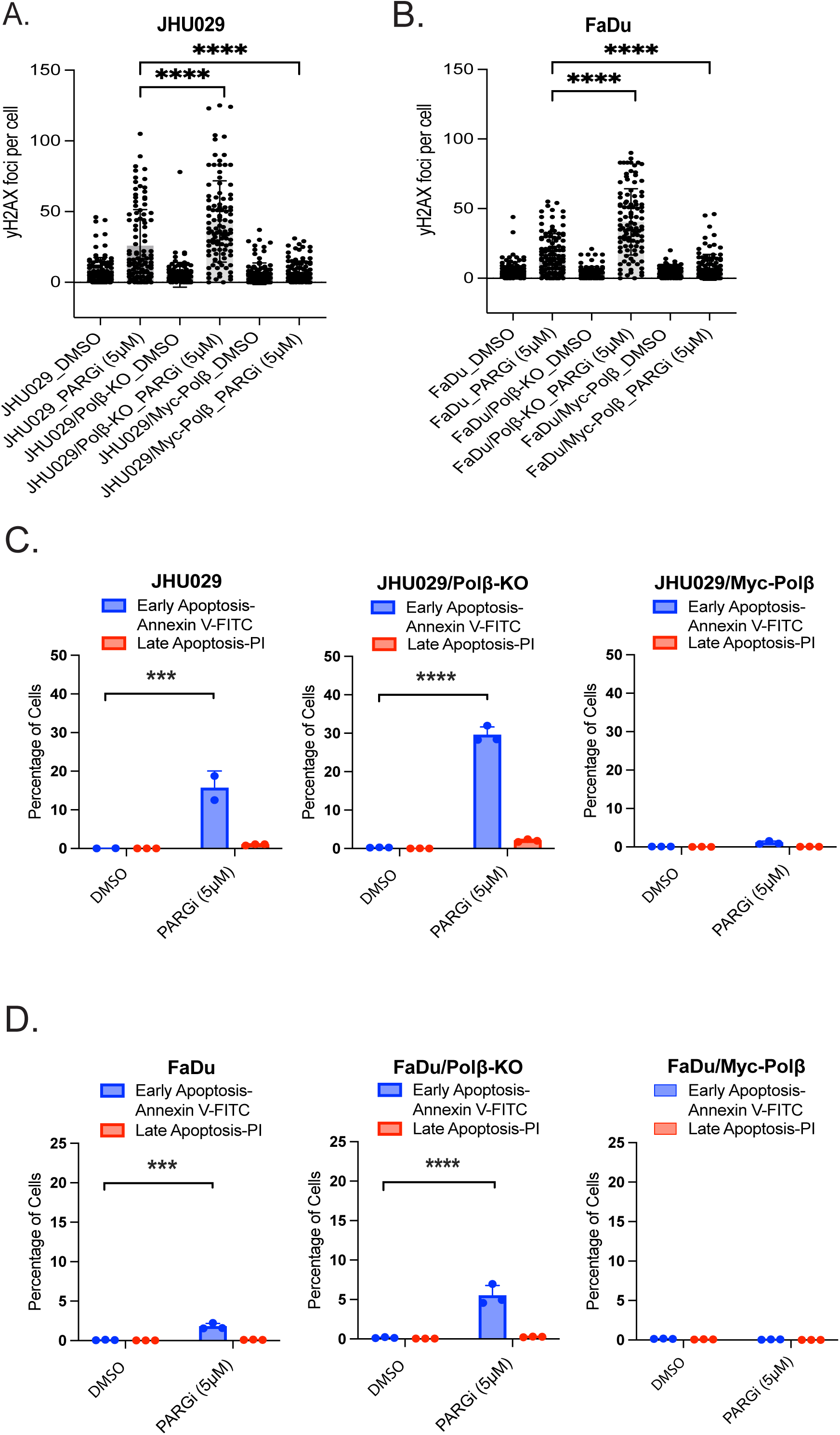
Effects of PARG Inhibitor on DNA damage and apoptosis in JHU029 and FaDu cells regulated by Polβ expression. A. γH2AX foci formation in JHU029, JHU029/Polβ-KO, and JHU029/Myc-Polβ cells following 24-hour treatment with PARG Inhibitor PDD00017273 (5 µM), with DMSO as control, n=3, >150 cells per each condition (p<****0.001; Mann-Whitney, two-tailed). B. γH2AX foci formation in FaDu, FaDu/Polβ-KO, and FaDu/Myc-Polβ cells following 24-hour treatment with PARG inhibitor PDD00017273 (5 µM), with DMSO as control, n=3, >150 cells per each condition, (p<****0.001; Mann-Whitney, two-tailed). C. Apoptosis assay evaluating early and late apoptosis in JHU029, JHU029/Polβ-KO (CRISPR/Cas9 knockout), and JHU029/Myc-Polβ cells treated with PARG inhibitor PDD00017273 (5 µM) for 72 hours, with DMSO as control, n=3 (the results are presented as mean ± SD. p*<0.05, p**<0.01, p***<0.001; Student’s t-test). D. Apoptosis assay evaluating early and late apoptosis in FaDu, FaDu/Polβ-KO, and FaDu/Myc-Polβ cells treated with PARG inhibitor PDD00017273 (5 µM) for 72 hours, with DMSO as control, n=3 (the results are presented as mean ± SD. p*<0.05, p**<0.01, p***<0.001; Student’s t-test).

Similarly, we assessed the sensitivity of FaDu, FaDu/Polβ-KO, and FaDu/Polβ overexpression cells to PARG inhibitors by evaluating γH2AX foci formation (**Figs. 2B, S1B**). FaDu cells displayed slight resistance to the PARG inhibitor, likely due to their functional DNA repair machinery, resulting in moderate γH2AX foci levels. In contrast, FaDu/Polβ-KO cells showed pronounced sensitivity, with the highest γH2AX foci formation, as the lack of Polβ impairs the BER pathway [6]. FaDu/Polβ overexpression cells exhibited increased resistance, forming the fewest γH2AX foci, owing to enhanced BER efficiency, which mitigates DNA damage induced by PARG inhibition.

### 3.4. Apoptosis analysis reveals Polβ-dependent PARG inhibitor resistance in HNSCC cells

Annexin V and PI staining were used to assess whether PARG inhibition induces apoptosis dependent on the level of Polβ expression in JHU029 and FaDu cells. We first conducted an apoptosis assay to evaluate the effect of PARG inhibitors on JHU029, JHU029/Polβ-KO, and JHU029/Polβ overexpression cells (**Figs. 2C, S1C**). JHU029 cells exhibited slight resistance to the PARG inhibitor, showing a low level of early apoptosis and low levels of late apoptosis. This indicates that while some cells undergo apoptosis in response to PARG inhibition, the inherent DNA repair capacity of the parental cells, facilitated by functional Polβ, allows for a significant proportion of cells to survive.

The absence of Polβ impairs the BER pathway [6], and inhibition of PARG in these cells leads to extensive accumulation of unrepaired DNA damage, triggering early apoptosis. This underscores the critical role of Polβ in DNA damage repair and cell survival. JHU029/Polβ-KO cells demonstrated heightened sensitivity to PARG inhibition, with the highest levels of early apoptosis and low levels of late apoptosis. Conversely, JHU029/Polβ overexpression cells showed increased resistance to PARG inhibition, with very low levels of early apoptosis and low levels of late apoptosis.

Similarly, apoptosis assays were performed to determine if the PARG inhibitor selectively induces apoptosis in FaDu parental cells, as compared to the FaDu/Polβ-KO, and FaDu Polβ-overexpressing cells (**Figs. 2D, S1D**). FaDu cells showed moderate early apoptosis and minimal late apoptosis, suggesting that their DNA repair capacity, supported by functional Polβ, mitigates PARG inhibitor-induced damage, allowing some cells to survive, like that seen for the JHU029 cells (**Fig. 2C**). Conversely, FaDu/Polβ-KO cells exhibited significant sensitivity, with the highest early apoptosis and minimal late apoptosis, due to the disruption of BER [6]. As with the JHU029, the FaDu/Polβ overexpression cells displayed enhanced resistance, with very low early and late apoptosis levels, highlighting the protective effect of efficient BER. The differential apoptotic responses underscore Polβ’s critical role in DNA repair and cell survival under PARG inhibition.

The differential apoptotic responses to PARG inhibition in the JHU029 and FaDu cell lines may be attributed to the role of Polβ in BER and a role in blocking the accumulation of replication-induced single-strand DNA gaps that have been suggested to modulate PARG inhibitor efficacy [19].

### 3.5. Overcoming treatment resistance with PARG and CHK1/ATR Inhibitors

Our earlier investigations demonstrated that the inhibition of PARG triggers a robust intra-S-phase response, activating the ATR and CHK1 signaling pathways [18, 21, 22]. The exploitation of PARG inhibition, combined with ATR/CHK1 inhibition, therefore, may influence cell viability, suggesting potential interactions between these two DNA damage response networks [35].

We used cell viability assays on JHU029 and FaDu cells to assess the cytotoxic potential of combined CHK1 and PARG inhibition. The cell-killing effects observed by the combination of a CHK1 inhibitor and a PARG inhibitor surpass the efficacy of the individual inhibitors, showcasing a promising strategy (**Fig. 3**). Cell viability was moderately reduced by treatment with either a PARG inhibitor (PARGi) or a CHK1 inhibitor (CHK1i), as illustrated in **Figs. 3A, 3B**). However, when PARGi and CHK1i were combined, the inhibition of cell growth was noticeably elevated and was found in both the FaDu and JHU029 cell lines.

**Fig. 3.**
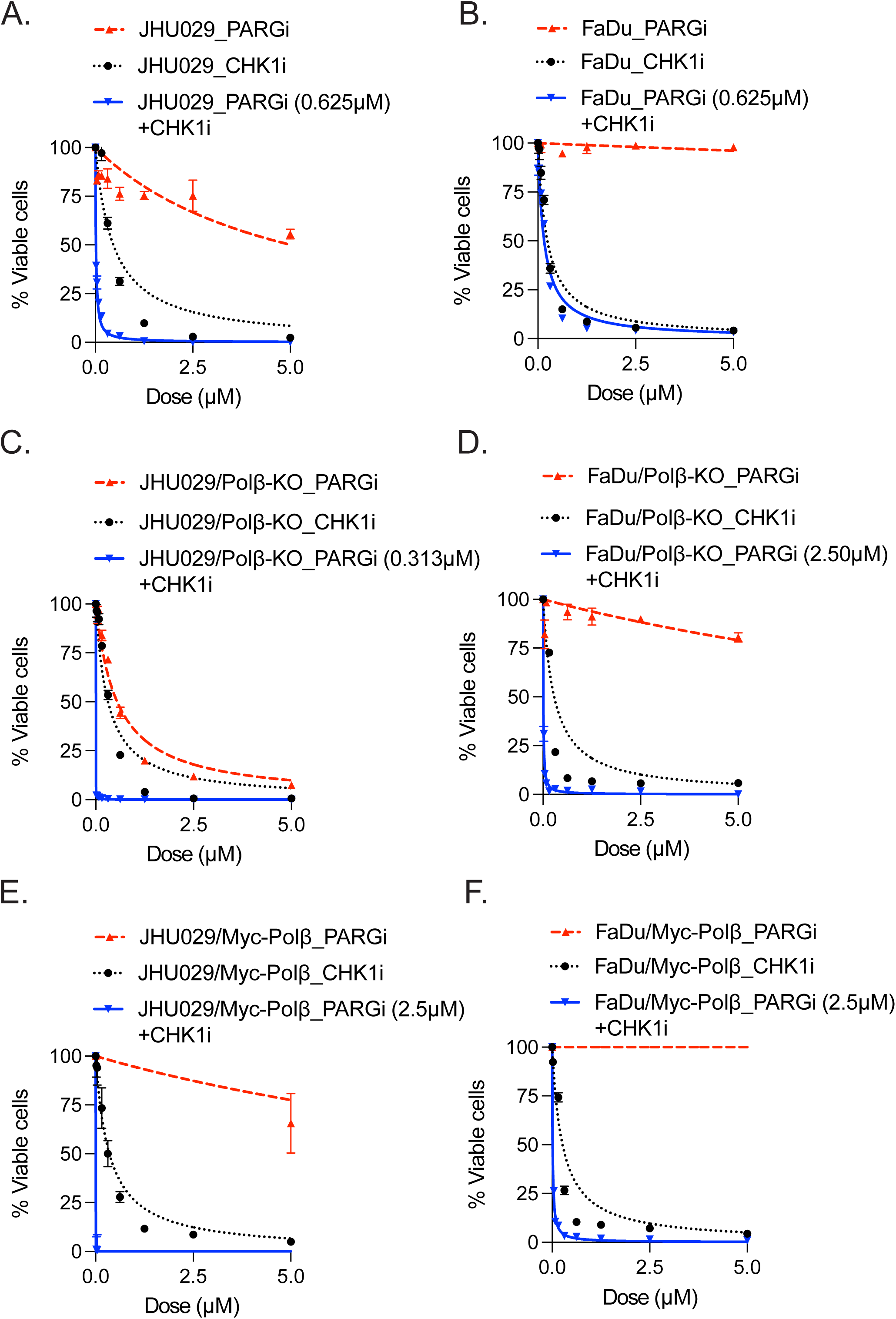
Cell viability of JHU029 and FaDu cells treated with CHK1 inhibitor alone or in combination with PARG inhibitor. A. JHU029 cell viability following treatment with PARGi (0.019 µM–5 µM), CHK1i (0.019 µM–5 µM) or PARGi (0.625 µM) + CHK1i (0.019 µM–5 µM) for 120 hours, as indicated, n=3 (comparison of PARGi, CHK1i or PARGi + CHK1i treatment at increasing dose: no stars = not significant, p*<0.05, p**<0.01, p***<0.001; Student’s t-test). B. FaDu cell viability following treatment with PARGi (0.019 µM–5 µM), CHK1i (0.019 µM–5 µM) or PARGi (0.625 µM) + CHK1i (0.019 µM–5 µM) for 120 hours, as indicated, n=3 (comparison of PARGi, CHK1i or PARGi + CHK1i treatment at increasing dose: no stars = not significant, p*<0.05, p**<0.01, p***<0.001; Student’s t-test). C. JHU029/Polβ-KO cell viability following treatment with PARGi (0.019 µM–5 µM), CHK1i (0.019 µM–5 µM) or PARGi (0.313 µM) + CHK1i (0.019 µM–5 µM) treatment for 120 hours, as indicated, n=3 (comparison of PARGi, CHK1i or PARGi +CHK1i treatment at increasing dose: no stars = not significant, p*<0.05, p**<0.01, p***<0.001; Student’s t-test). D. FaDu/Polβ-KO cell viability following treatment with PARGi (0.019 µM–5 µM), CHK1i (0.019 µM–5 µM) or PARGi (2.5 µM) +CHK1i (0.019 µM–5 µM) treatment for 120 hours, as indicated, n=3 (comparison of PARGi, CHK1i or PARGi +CHK1i treatment at increasing dose: no stars = not significant, p*<0.05, p**<0.01, p***<0.001; Student’s t-test). E. JHU029/Myc-Polβ cell viability following treatment with PARGi (0.019 µM–5 µM), CHK1i (0.019 µM–5 µM) or PARGi (2.5 µM) + CHK1i (0.019 µM–5 µM) treatment for 120 hours, as indicated, n=3 (comparison of PARGi, CHK1i or PARGi + CHK1i treatment at increasing dose: no stars = not significant, p*<0.05, p**<0.01, p***<0.001; Student’s t-test). F. FaDu/Myc-Polβ cell viability following treatment with PARGi (0.019 µM–5 µM), CHK1i (0.019 µM–5 µM) or PARGi (2.5 µM) + CHK1i (0.019 µM–5 µM) treatment for 120 hours, as indicated, n=3 (comparison of PARGi, CHK1i or PARGi + CHK1i treatment at increasing dose: no stars = not significant, p*<0.05, p**<0.01, p***<0.001; Student’s t-test).

Additionally, we extended this analysis to include cells lacking Polβ expression (JHU029/Polβ-KO, and FaDu/Polβ-KO) (**Figs. 3C, 3D**) and showed that, though to a smaller degree than in the parental lines, the combination of PARGi and CHK1i continued to exhibit a considerable decrease in cell viability. However, even though the elevated expression of Polβ (JHU029/Myc-Polβ and FaDu/Myc-Polβ cells) offered protection from the cell-killing effects of PARG inhibition alone, the combination of PARG and CHK1 inhibition could overcome Polβ-dependent resistance (**Figs. 3E, 3F**).

Similarly, we evaluated how cell viability was affected by ATR and/or PARG inhibition, alone, and in combination (**Fig. 4**). Cell viability was slightly decreased in JHU029 and FaDu cells treated with either PARGi or ATR inhibitor (ATRi), separately (**Figs. 4A, 4B**). However, there was a significant increase in cell killing when combined.

**Fig. 4.**
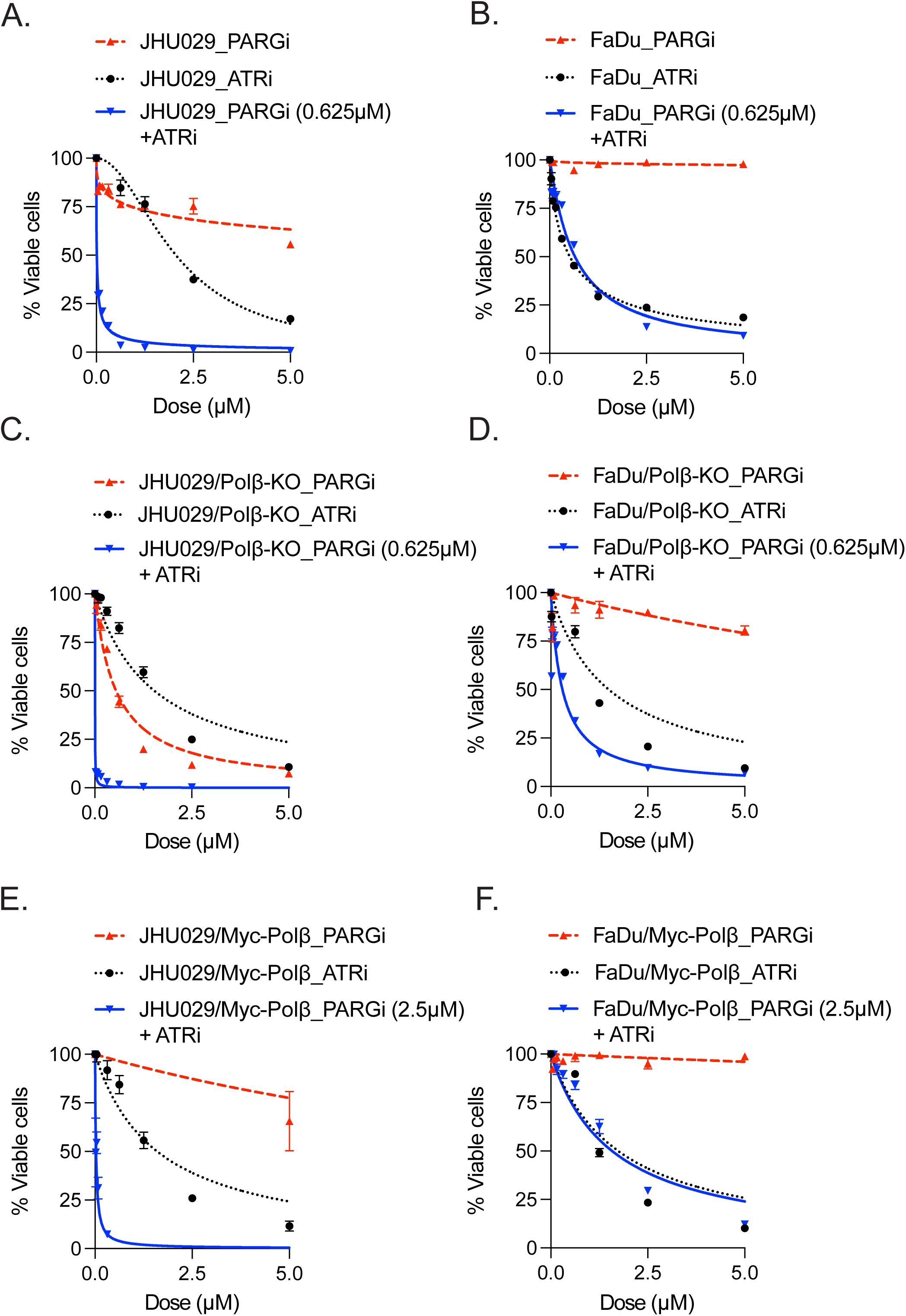
Cell viability of JHU029 and FaDu cells treated with ATR inhibitor alone or in combination with PARG inhibitor. A. JHU029 cell viability following treatment with PARGi (0.019 µM–5 µM), ATRi (0.019 µM–5 µM) or PARGi (0.625 µM) +ATRi (0.019 µM–5 µM) for 120 hours, as indicated, n=3 (comparison of PARGi, ATRi or PARGi +ATRi treatment at increasing dose: no stars = not significant, p*<0.05, p**<0.01, p***<0.001; Student’s t-test). B. FaDu cell viability following treatment with PARGi (0.019 µM–5 µM), ATRi (0.019 µM–5 µM) or PARGi (0.625 µM) +ATRi (0.019 µM–5 µM) for 120 hours, as indicated, n=3 (comparison of PARGi, ATRi or PARGi +ATRi treatment at increasing dose: no stars = not significant, p*<0.05, p**<0.01, p***<0.001; Student’s t-test). C. JHU029/Polβ-KO cell viability following treatment with PARGi (0.019 µM–5 µM), ATRi (0.019 µM–5 µM) or PARGi (0.625 µM) + ATRi (0.019 µM–5 µM) for 120 hours, as indicated, n=3 (comparison of PARGi, ATRi or PARGi + ATRi treatment at increasing dose: no stars = not significant, p*<0.05, p**<0.01, p***<0.001; Student’s t-test). D. FaDu/Polβ-KO cell viability following treatment with PARGi (0.019 µM–5 µM), ATRi (0.019 µM–5 µM) or PARGi (0.625 µM) + ATRi (0.019 µM–5 µM) for 120 hours, as indicated, n=3 (comparison of PARGi, ATRi or PARGi + ATRi treatment at increasing dose: no stars = not significant, p*<0.05, p**<0.01, p***<0.001; Student’s t-test). E. JHU029-Polβ/Myc-Polβ cell viability following treatment with PARGi (0.019 µM–5 µM), ATRi (0.019 µM–5 µM) or PARGi (2.5 µM) + ATRi (0.019 µM–5 µM) for 120 hours, as indicated, n=3 (comparison of PARGi, ATRi or PARGi + ATRi treatment at increasing dose: no stars = not significant, p*<0.05, p**<0.01, p***<0.001; Student’s t-test). F. FaDu-Polβ/Myc-Polβ cell viability following treatment with PARGi (0.019 µM–5 µM), ATRi (0.019 µM–5 µM) or PARGi (2.5 µM) + ATRi (0.019 µM–5 µM) for 120 hours, as indicated, n=3 (comparison of PARGi, ATRi or PARGi +A TRi treatment at increasing dose: no stars = not significant, p*<0.05, p**<0.01, p***<0.001; Student’s t-test).

In JHU029/Polβ-KO and FaDu/Polβ-KO cells, this trend continued (**Figs. 4C, 4D**), with the PARGi/ATRi combination showing a greater effect than with each inhibitor alone. Importantly, the combined treatment showed a significant decrease in viability of Polβ-overexpressing cells (JHU029/Myc-Polβ and FaDu/Myc-Polβ) (**Figs. 4E, 4F**), overcoming Polβ-dependent resistance. The potential of dual PARG and ATR inhibition is highlighted by these findings, especially in cells with increased Polβ expression.

## 4. Discussion

HNSCC represents a formidable challenge in oncology, with treatment resistance posing a significant barrier to effective therapy [1]. Current therapeutic approaches include radiation therapy and chemotherapy. Such DNA-damaging treatment options are often offset by enhanced DNA repair capacity in target cancer cells, such as seen in some cancer stem cells [36, 37]. DNA damage repair is a complex and highly regulated process that often involves PAR-dependent signaling. The inhibition of PARG leads to the accumulation of PAR chains within the cell, disrupting PAR-signaling networks and DNA damage repair processes, ultimately inducing cell death [8, 38].

The present study emphasizes how Polβ plays a crucial role in regulating the cellular response to inhibition of PARG in HNSCC cells. Our studies have specifically focused on evaluating how PARG inhibitors can be leveraged to overcome treatment resistance, particularly in scenarios where Polβ is overexpressed. Overexpression of Polβ, a DNA polymerase involved in the BER pathway, has been implicated in resistance to DNA-damaging therapies, posing a significant challenge in cancer treatment [39]. We have shown that overexpression of Polβ may also be related to genetic ancestry as we have previously reported AIMs linked to the altered mRNA expression of Polβ, with discernible elevation among African American patients compared to their European American counterparts that leads to a therapy-specific (radiation and/or platinum-based) poor survival response [4]. The mechanism linking overexpression of Polβ and this racial disparity in treatment response has not been clearly elucidated. However, our findings here provide some insight.

Employing molecular and cellular techniques, we elucidate the role of Polβ in modulating cellular responses to DNA damage and PARG inhibition. Our findings suggest that Polβ overexpression mitigates the cytotoxic effects of PARG inhibition. Additionally, Polβ overexpression alters apoptotic responses and cell cycle dynamics, further contributing to resistance mechanisms in HNSCC cells in response to PARG inhibition.

While Polβ overexpression significantly increases resistance to PARG inhibitors, loss of Polβ expression (knockout) makes cells susceptible to the cytotoxicity caused by PARG inhibition. This dynamic supports earlier research linking the effectiveness of BER to treatment resistance by highlighting Polβ as a critical determinant in DNA damage repair pathways [10, 30]. As we show herein, Polβ-overexpressing cells showed significant resistance to PARG inhibitors, even at higher concentrations, while Polβ knockout cells showed increased sensitivity. We show that Polβ reduces the cytotoxic accumulation of PAR at replication forks, likely by improving BER efficacy [6]. The reduced viability of Polβ-KO cells in response to PARG inhibition indicates the susceptibility of cells deficient in Polβ-mediated BER and implicates Polβ as a potential biomarker for predicting PARG inhibitor sensitivity.

The formation of PAR foci varied among cell lines after PARG inhibition, as shown by confocal microscopy. Polβ-KO cells had the most robust level of PAR foci. This would suggest that Polβ deficiency leads to elevated checkpoint activation and replication stress due to the accumulation of an elevated level of PARylated proteins. Conversely, the reduced level of PAR foci in cells overexpressing Polβ suggests a role for Polβ in maintaining replication progression and reducing replication-stress induced PAR signaling. These observations demonstrate the varied role of Polβ in regulating genomic integrity by regulating BER as well as reducing replication-associated stress.

Our results demonstrate differential responses to PARG inhibition among wild-type, Polβ-KO, and Polβ-overexpressing HNSCC cells. Polβ overexpression conferred significant resistance to PARG inhibitor-induced cell death, accompanied by reduced apoptotic responses. The cytotoxicity results were supported by apoptosis experiments, which showed that Polβ-KO cells had increased early apoptosis upon PARG inhibition, a sign of replication stress and unresolved DNA damage. Conversely, cells that overexpress Polβ showed decreased apoptotic responses, confirming Polβ’s protective function against cytotoxicity brought on by PARG inhibition. This finding supports the idea that Polβ increases BER, which promotes replication-associated lesion repair and inhibits apoptosis [40]. These results were further supported by analysis of γH2AX foci formation, which showed that Polβ-KO cells had the highest levels of γH2AX foci upon PARG inhibition, indicating significant DNA damage. Cells overexpressing Polβ showed fewer γH2AX foci, supporting the role of Polβ in reducing DNA damage and enhancing genome stability. Collectively, these findings demonstrate the significance of Polβ in preventing replication stress and preserving genomic integrity when PARG is inhibited.

This resistance to PARG inhibition in Polβ-overexpressing cells highlights the need for a combinatorial approach. Enhanced cytotoxicity, particularly in Polβ-overexpressing cells, is a significant finding of this study when PARG inhibitors are combined with ATR or CHK1 inhibitors. Despite their resistance to PARG inhibitors alone, Polβ-overexpressing HNSCC cells appear to be highly susceptible to dual pathway inhibition. The main regulators of the replication stress response are ATR and CHK1, which help in stabilizing and repairing replication forks in response to DNA damage [41]. By preventing PAR chains from degrading, PARG inhibition raises replication stress and prevents fork progression [18, 19, 21]. Cells overexpressing Polβ have higher BER activity, which inhibits the production of ssGAPs and prevents replication fork collapse [18]. However, this repair mechanism is disrupted when ATR or CHK1 are inhibited along with PARG, which results in enhanced cytotoxicity and ultimately leads to replication failure. We suggest that Polβ-overexpressing cells require ATR/CHK1 signaling to survive in the presence of increased PAR accumulation, highlighting the interaction of the BER and ATR/CHK1 pathways in reducing replication stress. This result is consistent with earlier research showing that by specifically targeting cells with increased replication stress, ATR and CHK1 inhibitors enhance the impact of drugs that damage DNA [42]. Targeting the ATR or CHK1 pathways, which are crucial for the replication stress response, could enhance the efficacy of PARG inhibitors in overcoming resistance [21]. The efficacy of ATR and CHK1 inhibitors also is found to require the hyper-activation of CDK2 [43] and so it will be interesting, in future studies, to determine if Polβ impacts CDK2 activation. Combining PARG inhibitors with ATR or CHK1 inhibitors may potentiate cell death in Polβ-overexpressing cancer cells by further compromising their ability to manage replication stress and DNA damage repair, offering a strategy to overcome PARG inhibitor resistance. The resistance conferred by Polβ overexpression highlights the need for novel approaches to overcome treatment challenges [14]. Our results suggest that combining PARG inhibitors with ATR or CHK1 inhibitors could offer a promising solution.

## 5. Conclusion

The significance of Polβ overexpression in improving BER efficiency and reducing replication stress is underscored by our findings in HNSCC cells. Polβ plays a critical role in regulating DNA repair, apoptosis, and cell cycle dynamics in response to PARG suppression, as indicated by analysis of wild-type, Polβ-KO, and Polβ-overexpressing cells. A promising approach to overcoming Polβ-mediated resistance may be the combination of PARG inhibitors and ATR or CHK1 inhibitors. This study establishes Polβ as a potential biomarker for predicting PARG inhibitor sensitivity and lays the groundwork for developing tailored approaches in HNSCC.

