## Supplementary Figures legends and Tables for "DNA polymerase beta expression in head & neck cancer modulates the poly(ADP-ribose)-mediated replication checkpoint"

### Supplemental Material

Supplemental Data include Supplemental Figures S1, S2 and Table S1.

### Figure Legends

**Supplemental Figure S1. Polβ-dependent effects of PARG inhibitor treatment on apoptosis and DNA damage in JHU029 and FaDu cells.**

A. γH2AX foci formation in JHU029, JHU029/Polβ-KO, and JHU029/Myc-Polβ cells following 24-Hour treatment with PARG Inhibitor PDD00017273 (5 μM), with DMSO as control.

B. γH2AX foci formation in FaDu, FaDu/Polβ-KO, and FaDu/Myc-Polβ cells following 24-Hour treatment with PARG Inhibitor PDD00017273 (5 μM), with DMSO as control.

C. Apoptosis assay evaluating early and late apoptosis in JHU029, JHU029/Polβ-KO, and JHU029/Myc-Polβ cells treated with PARG Inhibitor PDD00017273 (5 μM) for 72 hr, with DMSO as control.

D. Apoptosis assay evaluating early and late apoptosis in FaDu, FaDu/Polβ-KO, and FaDu/Myc-Polβ cells treated with PARG Inhibitor PDD00017273 (5 μM) for 72 hr, with DMSO as control.

Figure S1

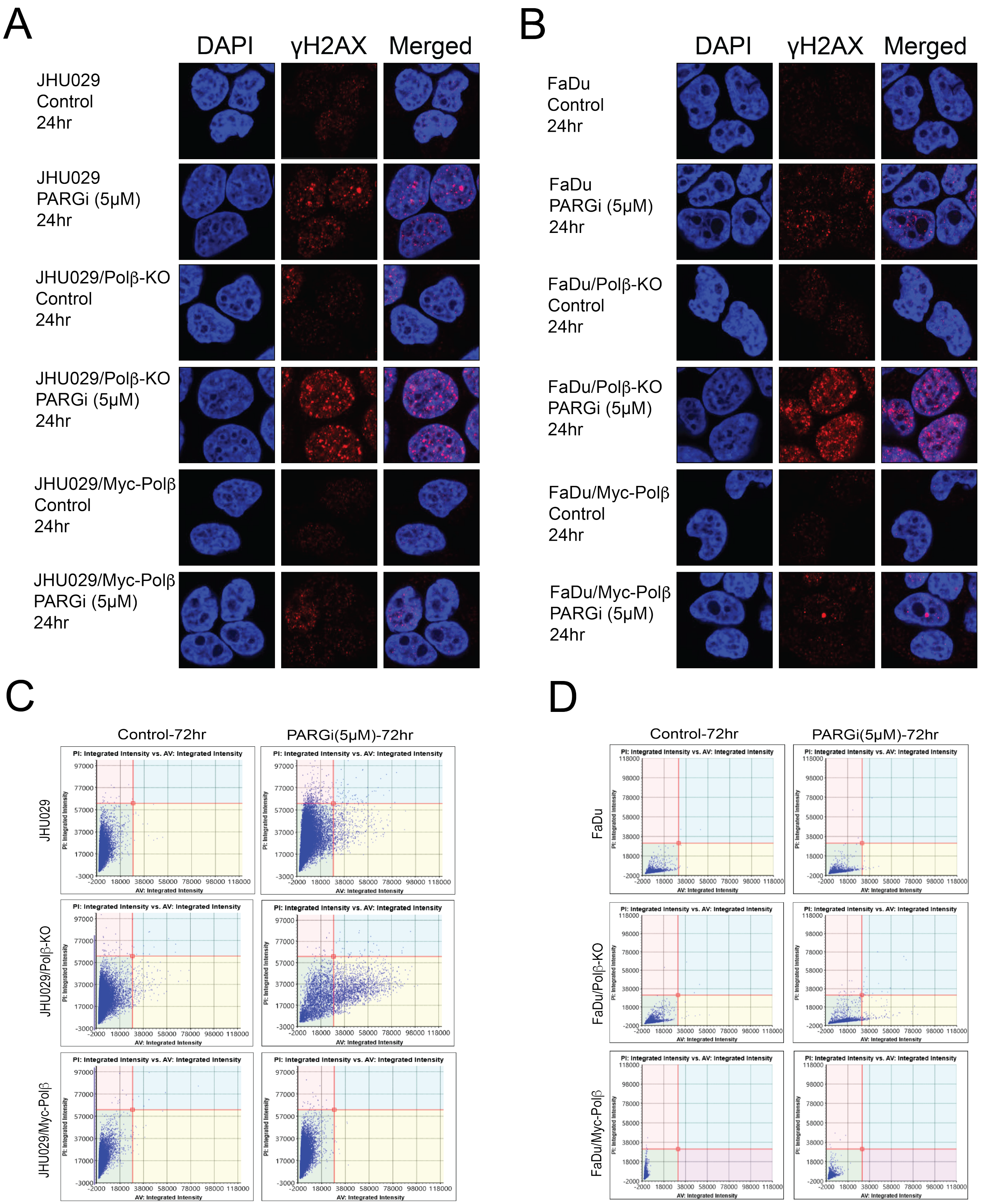
A
DAPI
γH2AX
Merged
JHU029 Control 24hr
JHU029 PARGi (5μM) 24hr
JHU029/Polβ-KO Control 24hr
JHU029/Polβ-KO PARGi (5μM) 24hr
JHU029/Myc-Polβ Control 24hr
JHU029/Myc-Polβ PARGi (5μM) 24hr
B
FaDu Control 24hr
FaDu PARGi (5μM) 24hr
FaDu/Polβ-KO Control 24hr
FaDu/Polβ-KO PARGi (5μM) 24hr
FaDu/Myc-Polβ Control 24hr
FaDu/Myc-Polβ PARGi (5μM) 24hr
C
Control-72hr
PARGi(5μM)-72hr
JHU029
JHU029/Polβ-KO
JHU029/Myc-Polβ
PI: Integrated Intensity vs. AV: Integrated Intensity
PI: Integrated Intensity
AV: Integrated Intensity
D
FaDu
FaDu/Polβ-KO
FaDu/Myc-Polβ

**Supplemental Figure S2. Full blots from immunoblots shown in Figure 1A.**

Shown are the full blots for the Polβ, XRCC1 and H3 immunoblots. The red box indicates the areas cropped to create **Figure 1A**.

Figure S2

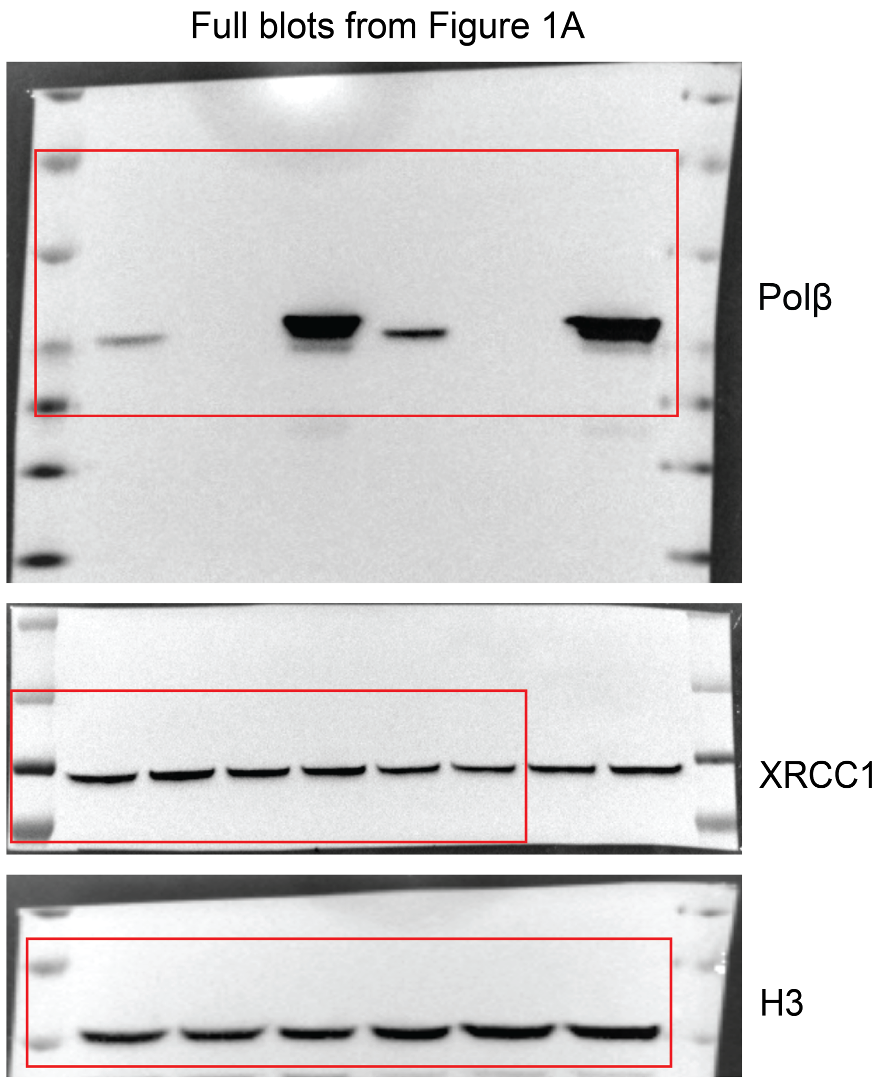

| **Table S1 - Reagents** | **Source** | **Identifier** |
|---|---|---|
| **Antibodies** (Use; Dilution) | | |
| Rabbit anti-XRCC1 (Immunoblot; 1:2000) | Bethyl Laboratories | Cat# A300-065A |
| Rabbit anti-Polβ (Immunoblot; 1:1000) | Abcam | Cat# ab175197 |
| Rabbit anti-H3 (Immunoblot; 1:5000) | Active Motif | Cat# 39451 |
| Rabbit anti-Phospho-Histone H2A.X (Ser139) (20E3) (Immunoblot; 1:1000) | Cell Signalling Technology | Cat# 9718S |
| Goat Anti-Rabbit IgG (H + L)-HRP Conjugate (Immunoblot; 1:2000) | Bio-Rad | Cat# 1706515 |
| Alexa Fluor 568 goat anti-rabbit antibody (1:500) | Invitrogen | Cat# A11011 |
| **Reagents, Chemicals and Peptides** | | |
| MEM | Gibco | Cat# 11095-080 |
| DMEM | Corning | Cat# 15-017-CV |
| RPMI 1640 | Gibco | Cat# 11875-119 |
| Heat-inactivated Fetal bovine serum | Atlanta Biologics | Cat# S11150 |
| Bovine serum albumin | RPI Research Products International | Cat# A30075–100.0 |
| L-glutamine | Gibco | Cat# 25030-081 |
| Penicillin/streptomycin | Gibco | Cat# 15140-122 |
| Antibiotic/antimycotic | Gibco | Cat# 15240-096 |
| Puromycin | Sigma-Aldrich | Cat# P9620-10ml |
| Hygromycin | Thermo Fisher Scientific | Cat# 10687010 |
| Trypsin-EDTA | Thermo Fisher Scientific | Cat# 25200-056 |
| Sodium Pyruvate | Invitrogen | Cat# 11360-070 |
| Non-Essential Amino Acids | Invitrogen | Cat# 11140-050 |
| 0.02μM Nitrocellulose membrane | Bio-Rad | Cat# 162-0115 |
| Protease inhibitor cocktail tablets | Thermo Fisher Scientific | Cat# 88666 |
| Blotting grade non-fat dry milk | Bio-Rad | Cat# 170-6404 |
| Nupage 4-12% Bis-Tris gel | Invitrogen | Cat# NP0323BOX |
| Clarity Western ECL Substrate | Bio-Rad | Cat# 1705060 |
| SuperSignal West Femto Maximum Sensitivity Substrate | Thermo Fisher Scientific | Cat# 34095 |
| Polybrene | Sigma-Aldrich | Cat# 107689 |
| 0.45μM Durapore Steriflip Filters | Sigma-Aldrich | Cat# SE1M003M00 |

| | | |
|---|---|---|
| Dimethyl Sulfoxide | Thermo Fisher Scientific | Cat# BP231-1 |
| CHK1 inhibitor (MK8776) | Selleckchem | Cat# S2735 |
| ATR inhibitor (Ceralasertib, AZD6738) | Selleckchem | Cat# S7048 |
| PARG inhibitor (PDD00017273) | Sigma-Aldrich | Cat# SML1781 |
| Hoechst 33342 | Thermo Fisher Scientific | Cat# 62249 |
| Paraformaldehyde, 4% in PBS | Thermo Fisher Scientific | Cat# J61899.AP |
| ViaStain No-Wash Annexin V-FITC Kit for Celigo | Revvity | Cat# CSK-V0007-1 |
| VECTASHIELD Antifade mounting medium with DAPI | Vector Laboratories | Cat# H-1200 |
| NucBlue Fixed Cell Stain Ready Probes | Thermo Fisher Scientific | Cat# R37606 |
| Cas9 Nuclease 3NLS | Synthego | |
| Opti-MEM™ I Reduced Serum Medium | Thermo Fisher Scientific | Cat# 31985062 |
| Lipofectamine CRISPRMAX Transfection Reagent | Thermo Fisher Scientific | Cat# CMAX00008 |
| RNAse | Thermo Fisher Scientific | Cat# EN0531 |
| **Cell lines** | | |
| 293-FT (derived from human embryonal kidney cells transformed with the SV40 large T antigen) | Thermo Fisher Scientific | Cat# R70007 |
| FaDu (Hypopharyngeal squamous cell carcinoma tumor cell line) | ATCC | Cat# HTB-43 |
| FaDu/Polβ-KO (FaDu cells transfected with purified Cas9 and three gRNA targeting distinct exon of POLB gene) | This study | |
| FaDu/Myc-POLB (FaDu cells expressing Myc fused to the N-terminus of PolB containing a mutation in the PAM site used by PolB gRNA1 & a hygromycin resistance cassette; Sobol Lab Stock 1861) | This study | |

| | | |
|---|---|---|
| JHU029<br>(Head and neck squamous cell carcinoma cells) | Gift from David Sidransky and Jeffrey N Myers | [1] |
| JHU029/Polβ-KO<br>(JHU029 cells transfected with purified Cas9 and three gRNA targeting exon 1 of the POLB gene) | This study | |
| JHU029/Myc-POLB<br>(JHU029 cells expressing Myc fused to the N-terminus of PolB containing a mutation in the PAM site used by PolB gRNA1 & a hygromycin resistance cassette; Sobol Lab Stock 1861) | This study | |
| **Oligonucleotides/gRNAs** | | |
| POLB multi gRNA<br>GGCCGCCAUGAGCAAACGGA<br>GUGCUAACCUGUGAGCAUGU<br>GUCGGUGUGGGAGAGAAGGA | This study | POLB-KO<br>(Synthego) |
| **Recombinant DNA** | | |
| pMDLg/pRRE | SLS# 253 | Addgene, Cat# 12251 |
| pRSV-Rev | SLS# 254 | Addgene, Cat# 12253 |
| pMD2.G | SLS# 252 | Addgene, Cat# 12259 |
| pLV-Hygro-EF1a-myc-PolB(PAMmut)<br>(Myc fused to the N-terminus of PolB containing a mutation in the PAM site used by PolB gRNA1 & a hygromycin resistance cassette) | [2],<br>SLS# 1861 | Addgene, Cat#176150 |
| pLV-EF1A-LivePAR-Hygro<br>(PAR binding domain with EGFP tag & a hygromycin resistance cassette) | [2-4],<br>SLS# 1727 | Addgene, Cat# 176063 |
| **Software and Algorithms** | | |
| Image J | Image J 1.48v | http://imagej.nih.gov/ijjava<br>1.6.0_65 |
| Adobe Illustrator (for preparation of figures) | Adobe Systems | Version 2024 |
| GraphPad Prism | GraphPad | Version 9.21<br>(Mac OS X) |
